# Nanopore-based genome-wide DNA methylation sequencing data of *Nosema ceranae*-inoculated and un-inoculated eastern honeybee workers’ midguts

**DOI:** 10.1101/2021.11.23.469791

**Authors:** Kejun Yu, Dafu Chen, Rui Guo

## Abstract

*Apis cerana cerana* is an excellent subspecies of *Apis cerana*, playing a vital role in pollination for wild flowers and crops as well as ecological balance. *Nosema ceranae*, an emergent fungal parasite infecting various bee species, originates from eastern honeybee. In this article, midguts of *N. ceranae*-inoculated *A. c. cerana* workers at 7 days post inoculation (dpi) and 10 dpi (AcT1 and AcT2) and un-inoculated workers’ midguts (AcCK1, AcCK2) were subjected to Nanopore-based genome-wide DNA methylation sequencing. Totally, 1773258, 2151476, 1927874 and 2109961 clean reads were generated from AcCK1, AcCK2, AcT1, and AcT2 groups, with the N50 lengths of 7548, 7936, 7678, and 7291 and the average quality value of 8.97, 8.95, 9.24, and 8.98, respectively. Among these, 93.85%, 94.49%, 88.69%, and 81.27% clean reads could be mapped to the reference genome of *A. c. cerana*. In the aforementioned four groups, 2149685, 2614513, 1637018 and 2726985 CHG sites were identified; the numbers of CHH sites were 9581990, 11801082, 7178559, and 12342423, whereas those of CpG sites were 14325356, 15703508, 14856284 and 13956849, respectively. Additionally, there were 36114, 118867, 30249, and 82984 6mA methylation sites respectively discovered. These data can be used for identifying differential 5mC methylation and 6mA methylation engaged in response of eastern honeybee workers to *N. ceranae* infestation, and for investigating the 5mC or 6mA methylation-mediated mechanism underlying host response.

**Value of the data:**

- The data provide enrichment for information about 5mC and 6mA methylation in eastern honeybees.
- Our data contributes to clarification of the epigenetic mechanism underlying eastern honeybee worker’s response to microsporidian infestation.
- This presented data offer novel insights into interaction between *Apis cerana* and *Nosema ceranae*.

## Data

In total, as shown in Table 1, 1773258, 2151476, 1927874 and 2109961 clean reads were respectively derived from AcCK1, AcCK2, AcT1 and AcT2 groups after quality control. The N50 lengths were 7548, 7936, 7678, and 7291, while the N90 lengths were 3784, 3969, 3485, and 3122, respectively (**Table 1)**. Additionally, the average quality value were 8.97, 8.95, 9.24, and 8.98, respectively (**Table 1**). Among these clean reads, 1664217, 2033008, 1709883, and 1714844 clean reads could be mapped to the reference genome of *A. c. cerana*, and the mapping ratios were 93.85%, 94.49%, 88.69%, and 81.27%, respectively (**Table 2**). As for 5mC methylation, 2149685, 2614513, 1637018, and 2726985 CHG sites were identified in AcCK1, AcCK2, AcT1 and AcT2 groups; the numbers of identified CHH sites were 9581990, 11801082, 7178559, and 12342423, while those of CpG sites were 14325356, 15703508, 14856284, and 13956849, respectively (**Table 3**). Moreover, there were 36114, 118867, 30249, and 82984 6mA methylation sites respectively discovered in AcCK1, AcCK2, AcT1 and AcT2 groups (**Table 3**).

**Table 1.** Overview of Nanopore-based genome-wide DNA methylation sequencing of un-inoculated and *N. ceranae*-inoculated *A. c. cerana* workers’ midguts.

| Sample | Number of sequence | Number of base | N50 Length | N90 length | Mean length | Maximum length | Mean quality |
| --- | --- | --- | --- | --- | --- | --- | --- |
| AcCK1 | 1773258 | 10567867182 | 7548 | 3784 | 5959 | 65272 | 8.97 |
| AcCK2 | 2151476 | 13190893032 | 7936 | 3969 | 6131 | 76893 | 8.95 |
| AcT1 | 1927874 | 11167595562 | 7678 | 3485 | 5792 | 77322 | 9.24 |
| AcT2 | 2109961 | 11494642421 | 7291 | 3122 | 5447 | 56791 | 8.98 |

**Table 2.** Summary of mapping of clean reads to the *A. c. cerana* reference genome

| Sample | All clean reads | Mapped clean reads | Unmapped clean reads | Mapping ratio |
| --- | --- | --- | --- | --- |
| AcCK1 | 1773258 | 1664217 | 109041 | 93.85% |
| AcCK2 | 2151476 | 2033008 | 118468 | 94.49% |
| AcT1 | 1927874 | 1709883 | 217991 | 88.69% |
| AcT2 | 2109961 | 1714844 | 395117 | 81.27% |

**Table 3.** Statistics of various DNA methylation sites

| Sample | Number of CHG | Number of CHH | Number of CpG | Number of 6mA |
| --- | --- | --- | --- | --- |
| AcCK1 | 2149685 | 9581990 | 14325356 | 36114 |
| AcCK2 | 2614513 | 11801082 | 15703508 | 118867 |
| AcT1 | 1637018 | 7178559 | 14856284 | 30249 |
| AcT2 | 2726985 | 12342423 | 13956849 | 82984 |

## 2. Experimental Design, Materials, and Methods

### 2.1 Fungal spore preparation

According to the developed method by Cornman et al. [1] in conjunction with some minor modifications [2], clean spores of *N. ceranae* were prepared under lab condition. (1) workers were kept in 20 C for 5 minutes, and then, midgut tissues were separated using sterile dissection tweezers, followed by homogenization in distilled water, filtration with four layers of sterile gauze, and three times of centrifugation at 8000 rpm for 5 minutes at 4 C. (2) After discarding the supernatant, the spores were further purified utilizing discontinuous density gradient solution composed of 5 mL each of 25%, 50%, 75% and 100% Percoll solution through centrifugation at 14000 rpm for 90 min at 4 °C. (3) The spores were seriously isolated using a sterile syringe and then centrifuged again on a Percoll discontinuous density gradient solution to gain clean spores. (4) A bit of purified spores were subjected to PCR detection using previously described primers [3] and confirmed to be mono-specific. (5) The concentration of fungal spores was measured using a CL kurt counter (JIMBIO, China). The spore suspension was freshly prepared before use.

### 2.2 Experimental inoculation, artificial rearing, and midgut sample preparation

The inoculation and rearing of *A. c. cerana* workers as well as midgut sample preparation were performed according to the established procedure in our lab [4, 5]. Briefly, (1) To offer newly emerged Nosema-free workers, combs of sealed brood from a healthy colony were kept in an incubator at 34 ± 0.5 °C, 50% RH; the emergent workers were seriously transferred into cages in groups (n=35) and kept in the incubator at 32 ± 0.5 °C, 50% RH. (2) A feeder filled with 4 mL sterile 50% (w/v) sucrose solutionwas inserted into each cage; the worker bees were fed adlibitum for 24 h and then starved for 2 h; each worker per group was immobilized and fed with 5 μL of 50% (w/v) sucrose solution containing 1×10^6^ *N. ceranae* spores; to ensure the complete ingestion of sucrose solution, theinoculated workers were isolated for 30 minutes in individual vials. Workers in control groups were fed with 50% (w/v) sucrose solution in an identical approach. The cages were carefully inspected every 24 h and any dead workers removed. (3) Six workers from treatment group or control group were sacrificed at 7 dpi and 10 dpi, followed by careful dissection of midgut tissues, which were instantly frozen in liquid nitrogen and then preserved at -80 °C. The treatment groups were termed as AcT1 and AcT2 groups, whereas the control groups were termed as AcCK1 and AcCK2 groups.

### 2.3 Genomic DNA isolation, library construction, and third-generation sequencing

Firstly, genomic DNA of AcCK1, AcCK2, AcT1, and AcT2 groups were respectively isolated wiith a Universal Genomic DNA Extraction Kit (TaKaRa, Tokyo, Japan) following the instructions, followed by determination of purity, concentration, and completeness using Nanodrop, Qubit, and agarose gel electrophoresis. Secondly, by using gtube, these genomic DNA were broken into segments with a size of approximately 8 kb. Thirdly, the SQK-LSK109 kit (Nanopore, London, the United Kingdom) was used for library construction. The steps included DNA damage repair, end repair, purification with magnetic beads, library quantification using Qubit. The constructed libraries were subjected to sequencing using Oxford Nanopore PromethION system.

### 2.4 Quality control and processing of sequencing data

The original electrical signals generated from Nanopore were data with FAST5 format, which were transformed to data with FASTQ format on the basis of basecall using Guppy software. Further, adaptors, reads shorter than 500 bp, and low-quality reads were filtered to gain high-quality clean reads, which were then aligned to the *A. c. cerana* reference genome (assembly ACSNU-2.0) using minimap2 software [6].

### 2.5 Genome-wide identification of 5mC and 6mA methylation sites

Nanopolish softwarewas used to identify CpG sites by mapping clean reads to the reference genome of *A. c. cerana* [7]. In addition, clean reads were subjected to re-squiggle using tombo software followed by exploration of CHH, CHG, and 6mA sites following alternative model[8].

## Acknowledgments

We gratefully acknowledge financial support of the Earmarked Fund for Modern Agro-industry Technology Research System (CARS-44-KXJ7), the Outstanding Scientific Research Manpower Fund of Fujian Agriculture and Forestry University (xjq201814), the Master Supervisor Team Fund of Fujian Agriculture and Forestry University (Rui Guo).

